# A trimeric autotransporter enhances biofilm cohesiveness in *Yersinia pseudotuberculosis* but not in *Yersinia pestis*

**DOI:** 10.1101/2020.03.31.019323

**Authors:** Joshua T. Calder, Nicholas D. Christman, Jessica M. Hawkins, David L. Erickson

## Abstract

Cohesion of biofilms made by *Yersinia pestis* and *Yersinia pseudotuberculosis* (*Yptb*) has been attributed solely to an extracellular polysaccharide matrix encoded by the *hms* genes (Hms-ECM). However, mutations in the *Yptb* BarA/UvrY/CsrB regulatory cascade enhance biofilm stability without dramatically increasing Hms-ECM production. We found that treatment with proteinase K enzyme effectively destabilized *Yptb csrB* mutant biofilms, suggesting that cell-cell interactions might be mediated by protein adhesins or extracellular matrix proteins. We identified an uncharacterized trimeric autotransporter lipoprotein (YPTB2394), repressed by *csrB*, which has been referred to as YadE. Biofilms made by a Δ*yadE* mutant strain were extremely sensitive to mechanical disruption. Overexpression of *yadE* in wild-type *Yptb* increased biofilm cohesion, similar to biofilms made by *csrB* or *uvrY* mutants. We found that the Rcs signaling cascade, which represses Hms-ECM production, activated expression of *yadE*. The *yadE* gene appears to be functional in *Yptb* but is a pseudogene in modern *Y. pes*tis strains. Expression of functional *yadE* in *Y. pestis* KIM6+ altered the production of Hms-ECM and weakened biofilms made by these bacteria. This suggests that although the YadE autotransporter protein increases *Yptb* biofilm stability, it may be incompatible with Hms-ECM production that is essential for *Y. pestis* biofilm production in fleas. Inactivation of *yadE* in *Y. pestis* may be another instance of selective gene loss in the evolution of flea-borne transmission by this species.

**IMPORTANCE:** The evolution of *Yersinia pestis* from its *Y. pseudotuberculosis* (*Yptb*) ancestor involved gene acquisition and gene losses, leading to differences in biofilm production. Characterizing the unique biofilm features of both species may provide better understanding of how each adapts to its specific niches. This study identifies a trimeric autotransporter YadE that promotes biofilm stability of *Yptb* but which has been inactivated in *Y. pestis*, likely because it is not compatible with Hms polysaccharide that is crucial for biofilms inside fleas. We also reveal that the Rcs signaling cascade, which represses Hms expression in *Y. pestis*, activates YadE in *Yptb*. The ability of *Yptb* to use polysaccharide or YadE protein for cell-cell adhesion may help it produce biofilms in different environments.

## INTRODUCTION

Environmental persistence, host interaction, and transmission of *Yersinia* depend on biofilms, which are tightly regulated by both transcriptional and post-transcriptional control mechanisms [1, 2]. Arguably, the best studied *Yersinia* biofilms are those made by *Y. pestis* while in the flea digestive tract that block the proventriculus and increase transmission to new hosts during flea feeding. These biofilms require the HmsHFRS proteins to produce and export a polysaccharide extracellular matrix of poly-ß-1,6-N-acetylglucosamine that is crucial in forming and maintaining bacterial cell-cell attachments [3, 4]. Without high levels of Hms-dependent extracellular matrix (Hms-ECM), the biofilms formed by *Y. pestis* while in fleas are not sufficiently cohesive to cause proventricular blockage. Adaptation to flea-borne transmission was precipitated in part by mutations that led to high levels of Hms-ECM compared to its *Y. pseudotuberculosis* (*Yptb*) ancestral lineage [5-7].

Among these mutations, modification of the Rcs regulatory system was especially important in enhancing Hms-ECM production. The Rcs signaling system includes the inner membrane kinase RcsC and the phosphorelay protein RcsD. RcsC phosphorylates itself when an inducing signal is present, and that phosphate is passed to RcsD and then to RcsB. Phosphorylated RcsB is a transcriptional regulator, binding to target promoters either as homodimers or as heterodimers with the auxiliary protein RcsA or other proteins [8]. Normally, the system is kept in the ‘off’ state by an inner membrane protein IgaA that blocks signaling through RcsD. When an appropriate activating signal is present, the RcsF lipoprotein sensor interacts with IgaA and relieves the inhibition [9]. RcsAB heterodimers negatively regulate *Yersinia* biofilms by binding to the promoters of the diguanylate cyclase genes *hmsT* and *hmsD*, as well as the *hmsHFRS* operon itself [10-12]. The *rcsA* gene is inactive in *Y. pestis* due to an insertion in the open reading frame, and restoring the function of this gene prevents biofilm formation and flea blockage [13].

Like many bacteria, exopolysaccharide production is positively regulated in *Yersinia* by cyclic di-GMP made by diguanylate cyclases [14, 15]. When RcsAB repression of *hmsT* and *hmsD* transcription is eliminated, the resultant high levels of the second messenger enhance Hms-ECM production, likely by activating the HmsRS inner membrane proteins [11, 16, 17]. Two phosphodiesterases that degrade cyclic-di-GMP are functional in *Yptb* but not in *Y. pestis*. A *Yptb* mutant strain wherein these genes (*rcsA* and both phosphodiesterases) are replaced with *Y. pestis* non-functional alleles can block fleas, but not to the same extent as *Y. pestis* [5]. This indicates that additional biofilm-related differences exist between the two species.

Several other regulatory influences on biofilm production in both *Yptb* and *Y. pestis* have been identified. We recently reported that the BarA/UvrY two-component system represses biofilms in *Yptb* by activating the CsrB small RNA [18]. Although *Yptb* mutants lacking *csrB* or *uvrY* make more cohesive biofilms than the wild type strain, their production of Hms-ECM does not approach that of *Y. pestis*. This suggested that *Yptb* biofilms may contain additional extracellular matrix components that are responsible for their cohesion. In this study, we investigated the basis for the increased cohesiveness of *csrB* mutant biofilms in an effort to identify novel *Yptb*-specific biofilm features. We found that *Yptb* biofilms have a significant protein component not present in *Y. pestis*. We focused on the uncharacterized YPTB2394 protein (YadE) which is repressed by *csrB*. This predicted lipoprotein is a part of the trimeric autotransporter family of proteins that function in bacterial adhesion to host surfaces or bacterial cell-cell attachments in biofilms [19-21]. Here, we show that production of YadE leads to *Yptb* biofilms that strongly resist disruption. Expression of *yadE* is activated by the Rcs system, in contrast to repression of Hms-ECM by RcsAB. Conversely, *yadE* is a pseudogene in *Y. pestis* and we demonstrate that *yadE* expression alters the production of Hms-ECM and weakens *Y. pestis* biofilms.

## RESULTS

### *Yptb* biofilm cohesion requires proteins

We previously demonstrated that mutations in the BarA/UvrY two-component regulatory system, or in the CsrB small RNA activated by UvrY, increase cohesion of *Yptb* strain IP32953 biofilms [18]. In that study, lectin staining of the *csrB* mutant strain indicated only moderately increased Hms-ECM on the surface of these bacteria. This suggested the possibility of additional structural components that are increased in the *csrB* mutant strain which assist in holding *Yptb* biofilms together. Protein adhesins and extracellular DNA are present in the biofilm extracellular matrices of numerous other bacteria. We therefore treated wild type and *csrB* mutant *Yptb* biofilms with DNase or proteinase K enzymes prior to testing their dispersal in mechanical disruption assays (Fig. 1). As expected, the *csrB* mutant formed more cohesive biofilms than the wild type strain. DNase did not significantly increase disruption of either wild type or *csrB* mutant biofilms compared with the control saline treatment. In contrast, proteinase K markedly increased the proportion of the *csrB* mutant strain biofilms that were dispersed. *Y. pestis* biofilms were very stable in these assays and were not affected by proteinase K or DNase treatment.

**Figure 1.**
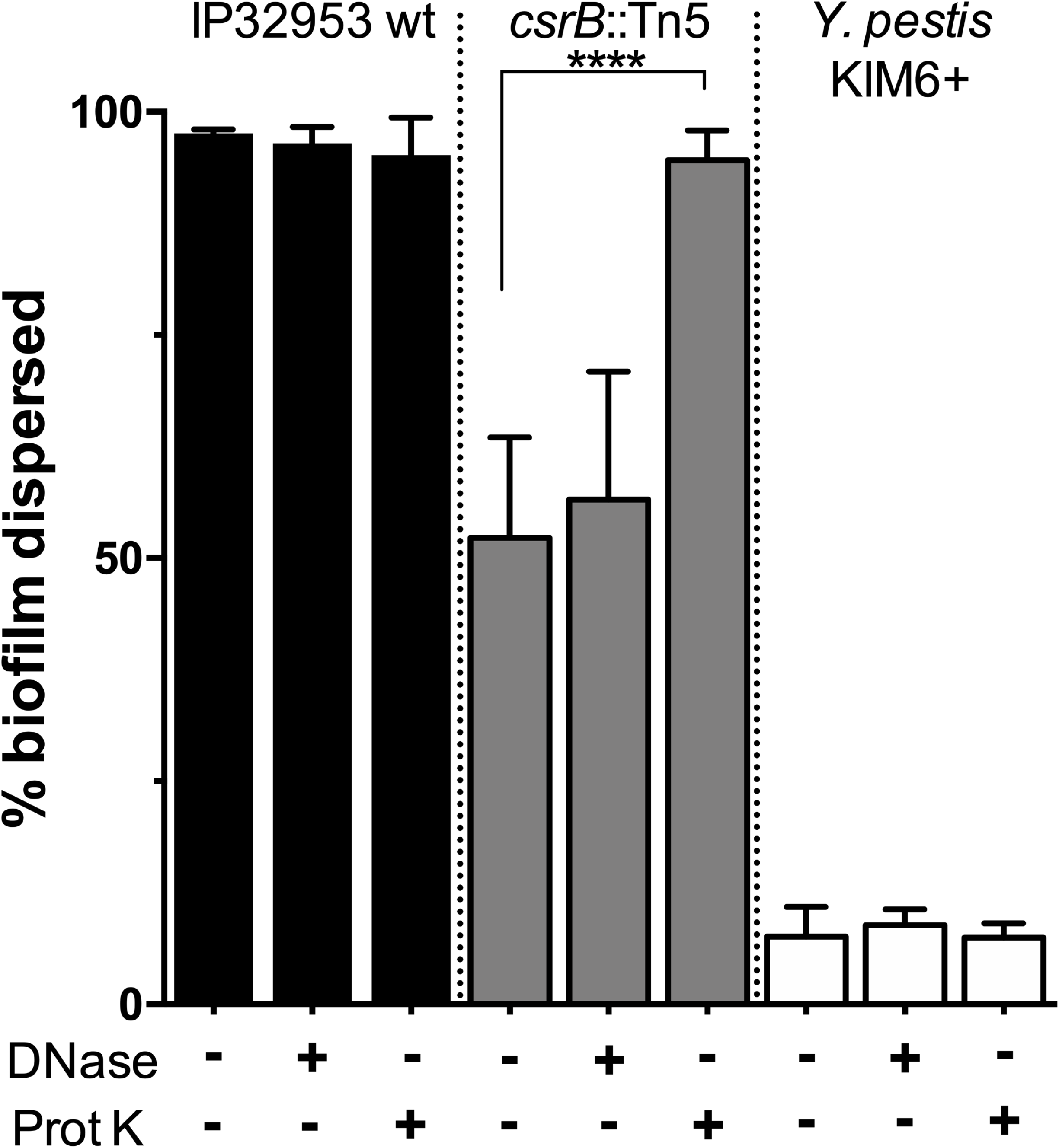
Increased stability of *Yptb csrB*::Tn5 mutant biofilms may be mediated by proteins. Biofilms were grown on polycarbonate filters for 72 hours. Filter biofilms were then treated with proteinase K or DNase for 30 minutes prior to agitation for 15 minutes. The proportion of the biofilms that were dislodged was measured by spectrophotometry. Pre-treatment of *Yptb csrB*::Tn5 mutant biofilms with proteinase K enhances their dispersal relative to the PBS control (****p<0.001 by Student’s T test) while DNase had no effect. *Y. pestis* biofilm stability was not affected by pretreatment with either enzyme.

### YPTB2394 putative autotransporter expression confers biofilm cohesion

Since we had demonstrated that *Yptb csrB* mutant biofilms are likely more cohesive in part due to a protein component of the extracellular matrix, we searched for proteins that were expressed more abundantly in the *csrB* mutant strain. We had previously conducted a comparison of the wild type and *csrB* mutant proteomes and found a large number of proteins that were expressed at lower levels in the *csrB* mutant, which is consistent with the role of CsrA as a translational repressor [15]. YPTB2394 was among the few proteins that were expressed more abundantly (approximately 30-fold higher) by the *csrB* mutant. YPTB2394 is annotated as a predicted lipoprotein with homology to type Vc autotransporter proteins. Trimeric autotransporter adhesins are membrane-anchored proteins known to mediate cell-cell attachments in other Gram-negative bacteria and promote greatly enhanced biofilm production in diverse species [22-24]. YPTB2394 is predicted to have a C-terminal YadA-like anchor, and several YadA-like stalk domains. The orthologous gene in *Y. pseudotuberculosis* strain YPIII (YPK_0761) has been referred to as *yadE* [25], although its function has not been investigated.

To examine the role of *yadE* in biofilm cohesion, we first deleted the gene from the wild type *Yptb* IP32953 strain. Biofilms formed by the mutant strain were extremely fragile compared to the wild type strain (Fig. 2). We then complemented the mutant strain by inserting *yadE* on a low-copy plasmid with a constitutive promoter. This resulted in biofilms that were highly resistant to disruption, even more so than *csrB*::Tn5 mutant biofilms. This strongly suggests that YadE helps maintain intracellular contacts and promotes *Yptb* biofilm cohesion.

**Figure 2.**
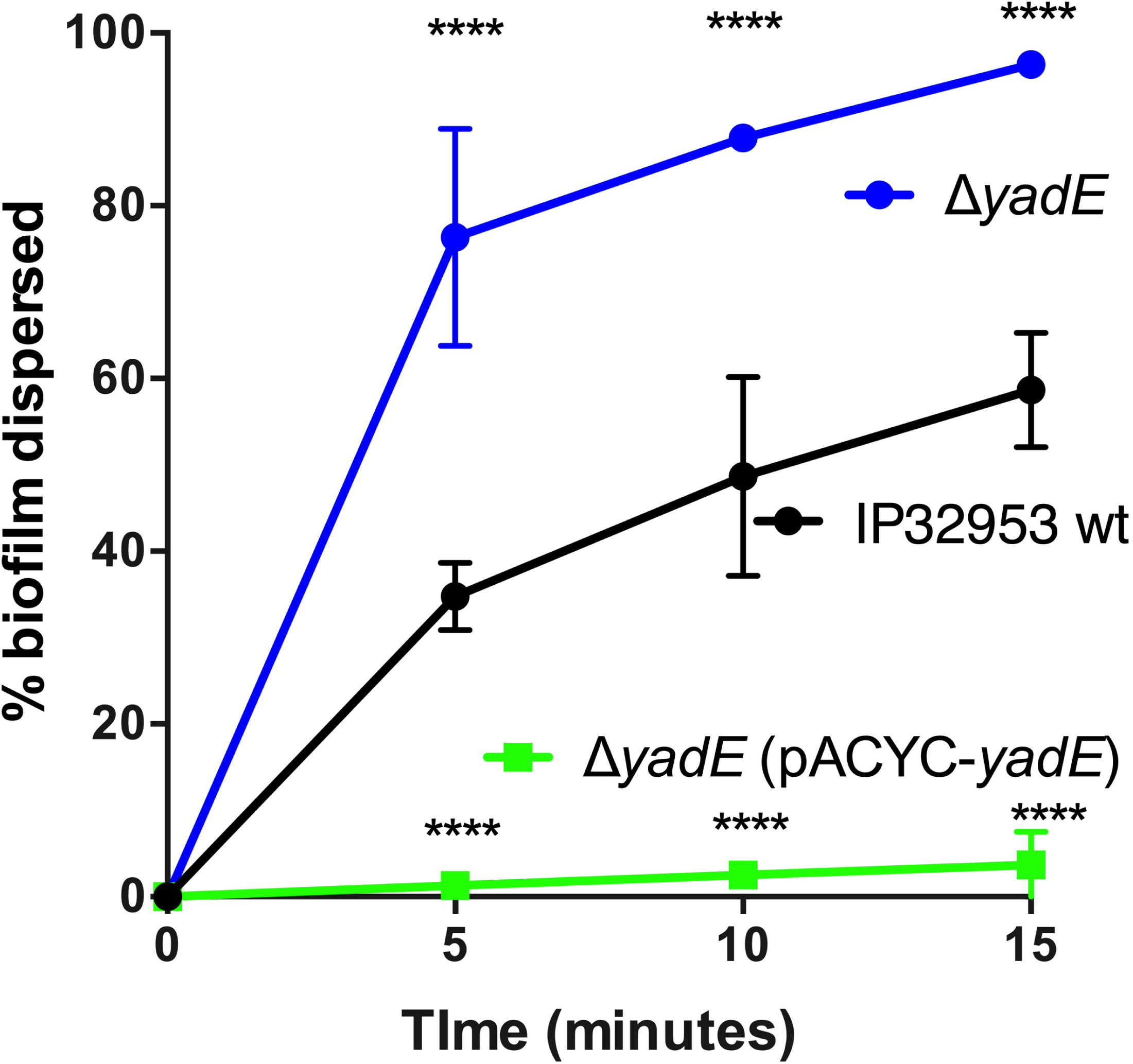
YPTB2394 (*yadE*) affects biofilm cohesion. Biofilms formed by the wild type IP32953, YPTB2394 (Δ*yadE*) mutant, and strain overexpressing *yadE* were each agitated and their dispersal measured at 5-minute intervals. Results were analyzed using repeated measures ANOVA with Tukey’s multiple comparison test (****significantly different from wild type, p<0.0001).

### *yadE* gene expression is regulated by CsrB and by the Rcs regulatory cascade

YadE (YPTB2394) protein levels are approximately 30-fold greater in *csrB* mutant bacteria compared to wild-type cells [18]. CsrB is a regulatory RNA that sequesters CsrA protein [26, 27]. Typically, CsrA binds to target sequences found near the Shine-Delgarno region of target transcripts and represses their translation. CsrB accumulation frees mRNA targets of CsrA to be translated more efficiently. However, translation of some CsrA-regulated proteins is enhanced by CsrA binding, and it is possible that CsrA is a direct translational activator of YadE. Alternatively, CsrA could repress translation of another transcriptional regulator, making its effect on *yadE* expression indirect. To further investigate regulation of *yadE*, we created a reporter plasmid containing the 356 bp region upstream of the start codon fused to a promoterless green fluorescent protein (*gfp*) gene. Transformation of wild type cells with this plasmid resulted in low but detectable levels of fluorescence. Expression was significantly enhanced in *csrB*::Tn5 mutant bacteria (Fig. 3), confirming the negative effect on YadE expression by CsrB.

**Figure 3.**
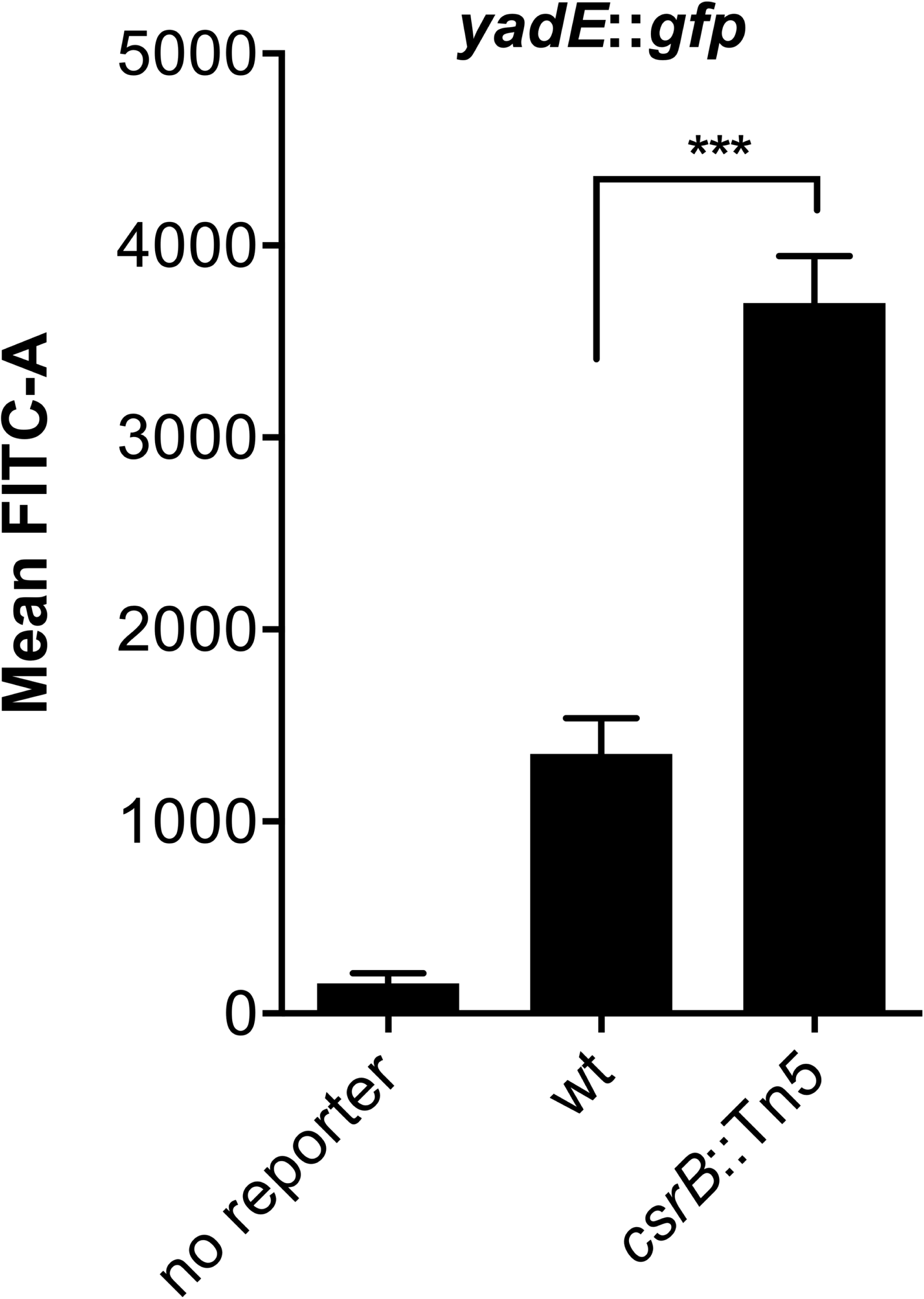
Expression of *yadE* is repressed by CsrB. The *Yptb* IP32953 wild type and *csrB*::Tn5 mutant strains were transformed with a *yadE*::*gfp* reporter plasmid. After growth for 48h on solid media the fluorescence of the population was measured by flow cytometry. *** indicates a significant difference from the wild-type strain (p=0.0002) by unpaired T-test.

In order to identify possible transcriptional repressors of *yadE*, we created a transposon mutant library in a wild type IP32953 strain carrying the *yadE*::*gfp* reporter plasmid. We then used fluorescence activated cell sorting to enrich for mutants with higher fluorescence than the wild type strain. After sorting, we verified enhanced *yadE::gfp* activity in isolated single colonies of 48 mutants by flow cytometry and determined their transposon insertion sites. We determined 23 distinct insertion sites among these mutants (Supplementary Table 1). We obtained 9 separate transposon mutants with insertions in the *igaA* gene, encoding the periplasmic IgaA protein that prevents overactivation of the Rcs signaling cascade [8]. We also obtained multiple mutants with insertions in genes encoding adenylate cyclase (*cya*) and the stringent starvation protein (*sspA*).. We tested biofilms made by *cya, sspA*, and *igaA* transposon mutants and found that they were also more cohesive than the wild type strain (Fig. 4), correlating with increased expression of *yadE*.

**Figure 4.**
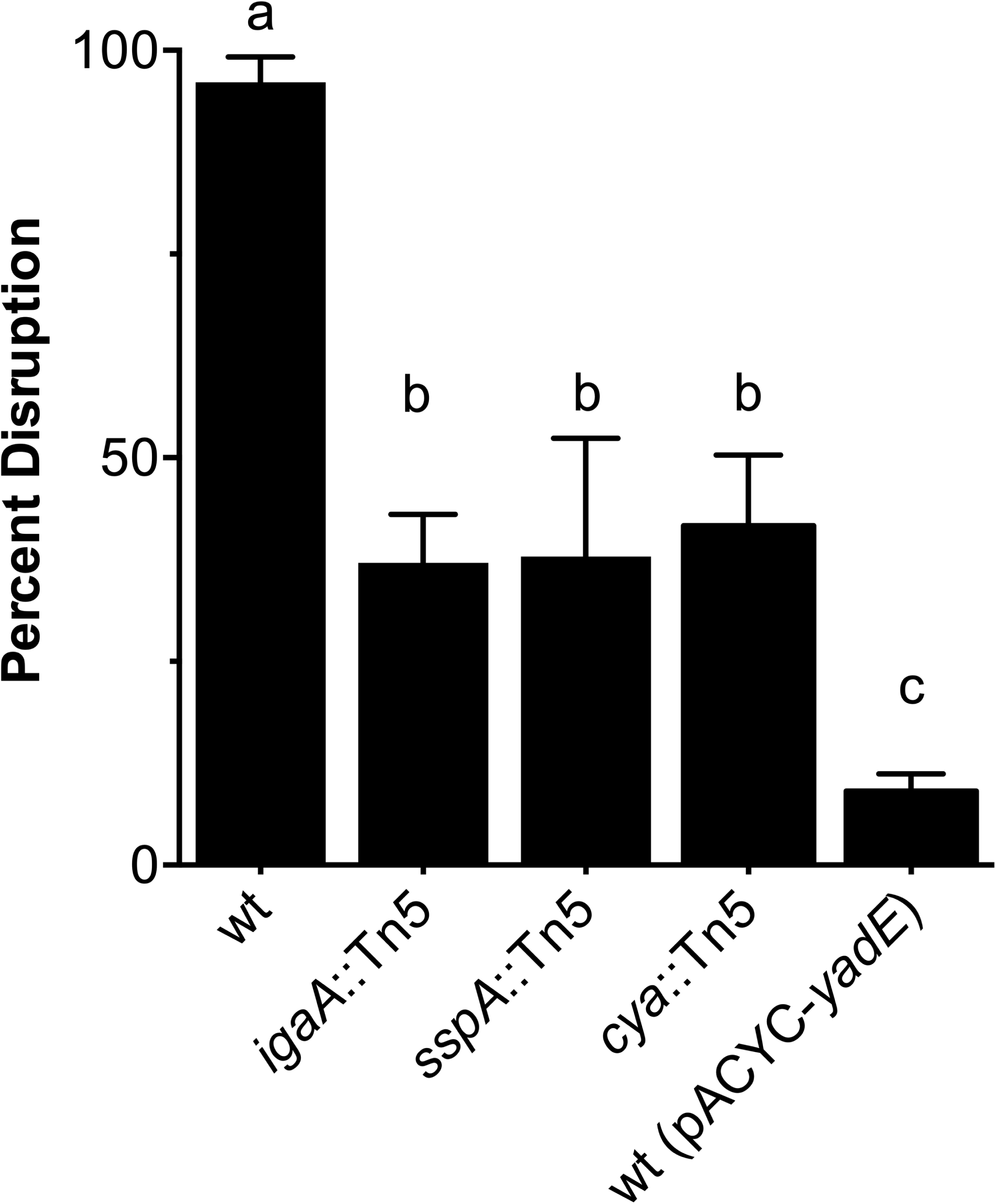
Mutants that overexpress *yadE* produce more cohesive biofilms. Transposon mutants derived from IP32953 with high *yadE*::*gfp* reporter activity were selected by FACS. Individual mutants with insertions in *igaA, sspA*, and *cya* genes were tested in biofilm disruption assays in comparison with the wild type IP32953 strain and the wild type strain overexpressing *yadE* (pACYC-*yadE*). One-way ANOVA analysis was performed with Tukey’s correction for multiple comparisons. Columns with the same letter (a, b, c) are not significantly different from each other (95% confidence interval).

IgaA limits the phosphorylation of RcsC and thereby prevents activation of RcsB. In some Enterobacteriaceae, *igaA* is an essential gene as overactivation of the Rcs cascade is lethal [28, 29]. In contrast, *Y. pestis* mutants with transposon insertions in *igaA* have been reported [30, 31] and our results confirm that this gene is also not essential in *Yptb* despite its fully functioning Rcs cascade. We verified that plasmid complementation of the *igaA* mutation could restore lower *yadE* expression similar to the wild type strain (Fig. 5). To confirm that overactive Rcs signaling was responsible for the enhanced *yadE* expression in *igaA* mutants, we compared *yadE* promoter activity in an *rcsB*::Tn5 mutant background, and in bacteria overexpressing *rcsB* via a multicopy plasmid. Lack of *rcsB* did not measurably decrease *yadE* expression compared to the wild type strain, but overexpression resulted in much higher fluorescence, similar to the *igaA* mutant (Fig. 5). These results indicate that the Rcs signaling cascade positively regulates *yadE* expression, opposite to Rcs repression of Hms-ECM production [11].

**Figure 5.**
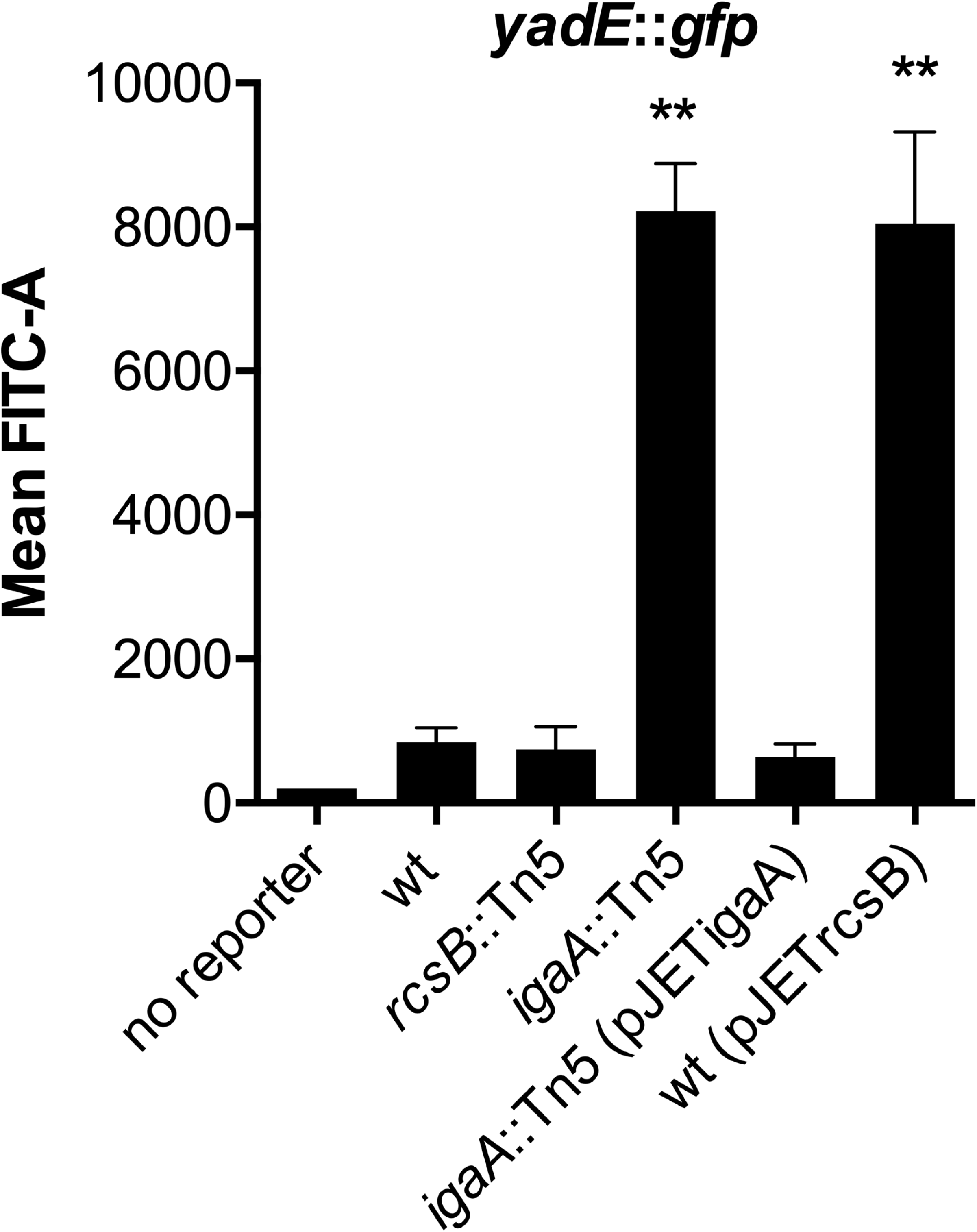
Expression of *yadE* is controlled by the Rcs phosphorelay. Inactivation of *igaA* or overexpression of *rcsB* increased *yadE*::*gfp* reporter activity. Fluorescence was measured by flow cytometry and compared with the wild type strain (** indicates a significant difference by unpaired T test, p<0.01).

### *yadE* is a pseudogene in *Y. pestis*, and expression of functional *yadE* prevents Hms-ECM production

Selective gene loss in *Y. pestis* during its divergence from *Yptb* has contributed to the emergence of flea-borne transmission [5, 6, 13, 32]. All publicly available genome sequences contain a *yadE* gene fragment that is identical among *Y. pestis* strains. Pairwise alignment shows four small deletions in the N-terminal region of the *Y. pestis* sequences relative to the *Yptb* IP32953 *yadE*. These are predicted to maintain the reading frame with 98.5% amino acid identity across the first 706 residues. However, a single insertion in a poly-G tract starting at nucleotide 2118 alters the reading frame of the *Y. pestis* sequence and results in five premature stop codons (Fig. 6). These are predicted to result in a non-functional protein missing the C-terminal YadA-like membrane anchoring domain. To investigate the possible consequences of *yadE* loss on *Y. pestis* biofilm stability, we transformed *Y. pestis* KIM6+ with the same plasmid conferring constitutive expression of the functional *yadE* gene. In contrast to its effect on *Yptb*, expression of *yadE* in *Y. pestis* resulted in biofilms that were more easily disrupted (Fig. 7a). This strain also produced colonies that were less pigmented on Congo-red agar plates, which is suggestive of reduced Hms-ECM production (Fig. 7b). We also tested the production of Hms-ECM using lectin staining and flow cytometry. Wild type *Y. pestis* binds strongly to wheat-germ agglutinin specific for N-acetylglucosamine polysaccharides. Conversely, *Y. pestis* expressing *yadE* exhibited a bipolar WGA-lectin staining pattern, where a significant portion of the cells did not bind to the lectin while another population bound at higher levels than the wild type strain (Fig. 7c). Thus, expression of functional *yadE* from *Yptb* in *Y. pestis* weakens biofilm cohesiveness, perhaps by altering production of the Hms-ECM.

**Figure 6.**
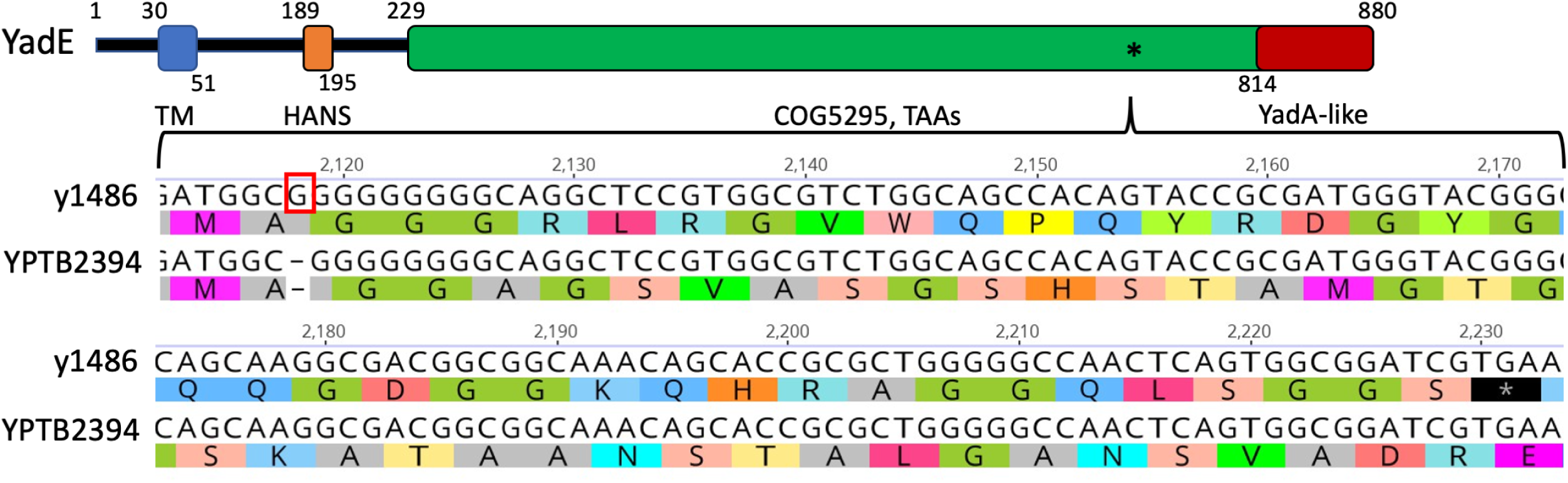
The YPTB2394 and y1486 orthologues of *Yptb* and *Y. pestis*. Domains predicted in the YPTB2394 protein sequence typical of trimeric autotransporters [61] include a transmembrane helix containing a secretion signal peptide (30-51), a HANS domain that typically connects α-to-β regions of proximal to head regions, COG5295 comprising the β-strand head domains with repetitive motifs, and the YadA-like anchor consisting of a 12-stranded outer membrane β-barrel. The asterisk notes the region containing the premature stop codons found in the *Y. pestis* sequences. Pairwise alignment of the *Yptb* IP32953 *yadE* (YPTB2394) and *Y. pestis* KIM pseudogene (y1486) nucleotide sequences shows that the sequences are highly similar across the 2643 bp *Yptb* sequence. All *Y. pestis* sequences have 14, 7, 4, and 5 bp deletions between nucleotides 185-209 and 326-340, which maintain the reading frame (not shown). A guanine insertion at nucleotide 2118 changes the reading frame and results in 5 predicted stop codons in the C-terminal region beginning at nucleotide 2230 that are predicted to eliminate the C-terminal YadA-like anchor.

**Figure 7.**
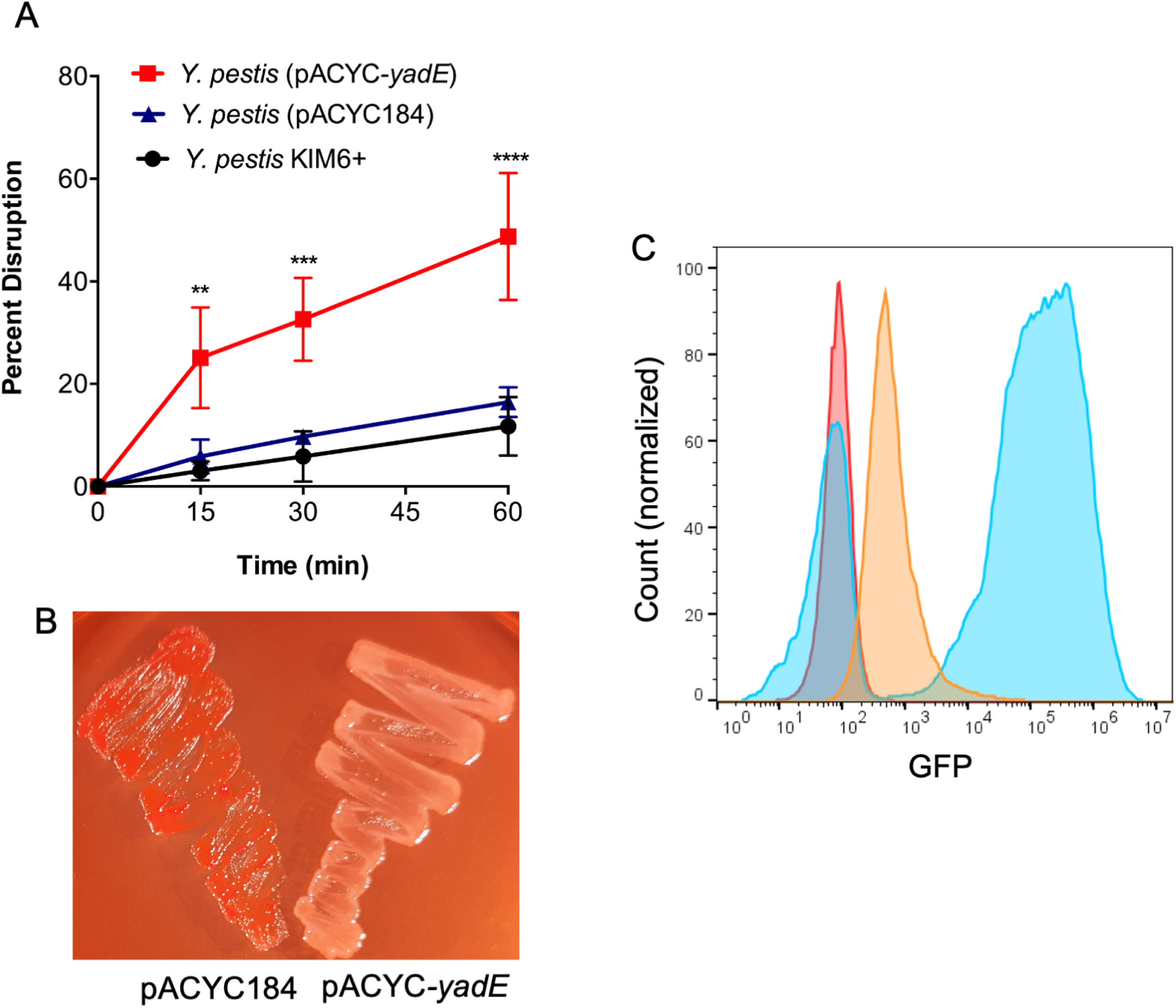
Expression of functional *yadE* in *Y. pestis* reduces biofilm stability and alters Hms-ECM production. A. Disruption of biofilms formed by *Y. pestis* KIM6+ wild type or strains transformed with pACYC-*yadE* or the pACYC184 empty vector. Results were analyzed using repeated measures ANOVA with Tukey’s multiple comparison test (**p<0.01, ***p<0.001, ****p<0.0001) B. Pigmentation of *Y. pestis* (*yadE*+ or empty vector) grown on Congo-red agar plates. C. Flow cytometry analysis of *Y. pestis* stained with GFP-labeled WGA lectin that binds to HMS-ECM. The wild type strain (orange) is uniformly labeled compared to the unstained control (red) while the strain expressing functional *yadE* (blue) exhibits a bipolar staining pattern with either no detectable lectin bound to the surface or with very high staining.

## DISCUSSION

Biofilms may be held together by extracellular polysaccharides, DNA, or proteins. Among *Yptb* strains, carriage of the *hms* genes encoding for polysaccharide extracellular matrix is ubiquitous, although they vary greatly in their expression of these genes [33]. The Hms-ECM is required for *Yptb* biofilm production in some conditions, including on the mouthparts of predatory nematodes [34, 35]. Extracellular DNA has also been detected in at least some *Yptb* nematode-associated biofilms, although the source and functional significance of the DNA remain unclear [36]. The extracellular matrix of biofilms formed by *Y. pestis* while they are in the flea digestive tract incorporates material derived from blood digestion [4], but the bacteria are not known to produce any other biofilm extracellular matrix material other than Hms-ECM. Our study is the first to demonstrate a role for a protein in mediating cohesion of *Yersinia* biofilms. The YPTB2394 (*yadE*) gene encodes a trimeric autotransporter protein vital to the stability of *Yptb* biofilms, as its inactivation results in a fragile biofilm and its overexpression significantly strengthens cohesiveness.

Multiple autotransporter proteins belonging to either type Va, Vc, or Ve families are present in *Y. pestis* and/or *Yptb* [37]. Type Va *Yersinia* autotransporters have been mainly investigated in the context of *Y. pestis* infection. YapE, YapJ, YapK, and YapV promote development of bubonic and/or pneumonic plague in mice [38-40], likely by enhancing adherence to epithelial cells. YapC can induce aggregation when expressed in *E. coli* but no effects on *Y. pestis* biofilms following deletion of this gene were observed [41]. *Yptb* strains also possess an additional type Va autotransporter gene *yapX*, whose function is unknown, which is also a pseudogene in all *Y. pestis* strains [38].

YadE is predicted to be a member of the type Vc trimeric autotransporter family. Numerous species use trimeric autotransporters to adhere to other bacteria within biofilms, host extracellular matrix proteins, or inanimate surfaces [21]. The prototype of the Vc family is YadA, made by *Yptb* and *Y. enterocolitica* but which has been inactivated in *Y. pestis*. YadA promotes autoagglutination and tight adherence to eukaryotic cells necessary for proper injection of effector proteins via type III secretion [42]. Restoring *yadA* gene function in *Y. pestis* has been reported to decrease its virulence in mouse infections [43]. It is not known whether YadA expression also interferes with Hms-ECM production by *Y. pestis*. However, it is co-expressed along with the type III secretion effectors encoded on the virulence plasmid at 37°C rather than at lower temperatures when Hms-ECM is produced. *Yptb* and *Y. pestis* both express two additional type Vc proteins, YadB and YadC, also at 37°C, which promote invasion of host cells and bacterial survival in skin [44].

Versatile biofilm production strategies are probably most helpful to bacteria such as *Yptb* that are found in many different environments (free-living or within amoeba in soil or water, in plants, or in the digestive tracts of multiple animals) [45]. Conversely, niche specialization might be promoted by the selection of one biofilm pathway at the expense of others. Many *Staphylococcus* strains produce biofilms that are dependent on polysaccharide intracellular adhesin, which is identical to Hms-ECM [46]. However, expression of this polymer tends to be suppressed in methicillin-resistant *S. aureus* strains in favor of fibronectin binding proteins [47]. Staphylococci that predominantly inhabit environments with high shear stress or grow on medical devices produce primarily polysaccharide-dependent biofilms [48, 49], whereas those that interact more directly with host tissues may benefit from protein-based biofilms that allow them to incorporate fibrin or other host proteins into a protective shield [50]. It is tempting to speculate that *Yptb* strains retain multiple biofilm strategies that provide flexibility according to changing environments, whereas *Y. pestis* may have jettisoned alternative biofilm strategies to Hms-ECM as it adapted to its restricted lifestyle of flea-rodent transmission.

Although *yadE* (y1486 in the *Y. pestis* KIM sequence) is a pseudogene in *Y. pestis*, it is one of the 100 most highly transcribed genes by this strain during infection of fleas [51]. Furthermore, even greater expression was measured in a *Yptb* mutant strain that is able to infect and block fleas [52]. Thus, the regulatory influences necessary for strong induction of this gene exist during flea infections. We found that Tn5 insertions in the adenylate cyclase responsible for producing cyclic AMP greatly enhance *yadE* expression. This is consistent with previously reported transcriptome comparisons of a cAMP receptor protein (CRP) mutant to the wild type YPIII strain [25]. Our results also demonstrate that induction of Rcs signaling, either due to inactivation of *igaA* or overexpression of *rcsB*, dramatically increases expression of *yadE*. The Rcs cascade is very complex and can respond to many different signals, including lipopolysaccharide and peptidoglycan perturbations [53, 54]. Innate immune defenses, osmotic changes, or other factors present in the flea digestive tract may inflict similar stresses to *Yersinia*. At the same time, high levels of *yadE* expression are disruptive to production, accumulation, or stability of the Hms-ECM. Hms-ECM is essential for proper biofilm formation and proventricular blockage, the major transmission mode of *Y. pestis* by rat fleas. Therefore, loss of *yadE* gene function may have been an additional key step in the divergence of *Y. pestis* from *Yptb*.

## EXPERIMENTAL PROCEDURES

### Bacterial strains, media and growth conditions

*Y. pestis* KIM6+ and *Y. pseudotuberculosis* IP32953 were grown at 21°C in Luria-Bertani (LB) or heart infusion broth containing 0.2% galactose (HIG) [18]. *E. coli* strains were grown in LB agar or broth at 37°C. Where required, kanamycin (30 µg/ml), chloramphenicol (10 µg/ml), or ampicillin (100 µg/ml) were added to media.

Deletion of *yadE* from *Yptb* IP32953 was accomplished through allelic exchange. The PCR primers used for creating the strains and plasmids for these experiments are listed in Supplementary Table S2. Upstream and downstream regions adjacent to YPTB2394 were cloned by overlapping extension PCR [55] and inserted into plasmid pRE112 [56] to create pREΔ*yadE.* The ampicillin resistance (*bla*) gene from pHSG415 [57] was then inserted into this plasmid replacing the *yadE* coding region to create pREΔ*yadE::bla*. After biparental mating with *E. coli* strain MFDλpir (pREΔ*yadE::bla*) [58] as donor and *Yptb* IP32953 as recipient, double-crossover mutations were selected on media containing 10% sucrose and ampicillin. The mutation was verified using PCR and Sanger sequencing (Eton Biosciences Inc.).

To overexpress *yadE*, the YPTB2394 gene as well as the upstream 200 bp region were placed under the control of the tetracycline resistance promoter on plasmid pACYC184. PCR products for the *yadE* gene and the pACYC184 backbone were combined in an overlapping-extension PCR reaction and transformed into *E. coli* DH5α to create plasmid pACYC-*yadE*. In a similar way, the pACYC-*yadE::gfp* reporter plasmid was created by amplifying the *gfp* coding region from pUC18R6K-miniTn7T sig70c35_GFP [59], the YPTB2394 promoter region, and the pACYC184 backbone. These PCR products were purified and combined in an overlapping-extension PCR reaction and transformed into *E. coli* DH5α. Plasmids containing *igaA* and *rcsB* (pJET-*igaA* and pJET-*rcsB*) were created by PCR amplification of the respective genes and cloning into the pJET1.2 plasmid (ThermoFisher). All plasmids were sequence verified and transformed into *Yersinia* strains by electroporation.

### Biofilm disruption assay

Biofilm stability was measured as previously described [18]. Overnight broth cultures of strains to be tested were transferred to polycarbonate track etched (PCTE) membrane filters (n=4-5) on HIG agar plates. After 72h of growth at 21°C, individual filters containing the biofilm samples were placed in 10 mL of phosphate buffered saline (PBS). Tubes were shaken vertically at 200 rpm and the optical density of dislodged cells was measured (A_600nm_) at specific time points. The filters were then vortexed at high speed until biofilms were completely disrupted and the A_600nm_ measured. Enzymatic treatment of biofilms prior to disruption tests were performed with proteinase K (Sigma) or DNase (Ambion) at 5 mg/ml. Solutions of enzyme in PBS (100 µl) were placed directly on top of the biofilms and incubated at 21°C for 60 minutes.

### Transposon mutagenesis and fluorescence activated cell sorting

A Tn5 transposon mutant library in strain IP32953 (pACYC-*yadE::gfp*) was created using the pRL27 donor plasmid as previously described [60]. After plating the mating mixture on HIB agar containing kanamycin and chloramphenicol, ∼100,000 individual mutant colonies were suspended and washed in PBS and diluted to 10^6^ cfu/ml for FACS sorting. A total of ∼4×10^6^ individual *Yptb* mutant bacteria were sorted using FACS Aria Fusion Cell Sorter (BD Biosciences) at the BYU Cell Sorting/Bio-Mass Spectrometry core facility. Cells exhibiting one standard deviation greater than the wild-type IP32953 strain in the GFP channel were collected, diluted and plated on HIB agar containing kanamycin and chloramphenicol to grow single colonies. The plates were then incubated for 24 h. Approximately 1000 colonies from these plates were examined individually using an LED mini-blue transilluminator (IO Rodeo). From this pool, 48 colonies were confirmed visually to express high levels of GFP after re-growth and were selected for further verification using flow cytometry. The Tn5 insertion sites in these mutants were determined by arbitrary PCR and sequencing as previously described [58]. Of the 48 colonies, 23 were found to have unique insertion sites as reported in Supplementary Table 1.

### Flow cytometry

To measure *yadE::gfp* expression, bacteria containing the reporter plasmid were grown on HIG agar plates at 21°C for 24 h. For each strain, bacteria from individual colonies were suspended in PBS and the GFP fluorescence measured by flow cytometry. For Hms-ECM measurement on the surface of bacteria, fixed bacterial cells were incubated with FITC-labeled wheat-germ agglutinin (Sigma) as previously described [18]. GFP expression and lectin binding of individual cells was measured using a BD Accuri C6 Flow Cytometer and analysed using FACSDiva software (BD Biosciences).

### Statistical analysis

Statistical analysis was performed using Graphpad Prism 6.0. The details for each test are provided in the relevant figure legend.

## ACKNOWLEDGEMENTS

We thank Daniel Call for technical assistance with cell sorting. JTC was funded through a BYU CURA grant.

---

**Supplementary Table 1.** Tn5 insertion mutants with increased *yadE::gfp* expression.

| **Tn5 Insertion point (orientation)** | **Mean fluorescence** | **YPTB gene number** | **Annotation** |
| --- | --- | --- | --- |
| Wild type IP32953 | 1352 |  |  |
| 70558 (-) | 1744 | YPTB0058 | *yibQ* hypothetical protein |
| 224414 (+) | 3395 | YPTB0185 | *cyaA* adenylate cyclase |
| 223538 (+) | 3104 | YPTB0185 | *cyaA* |
| 757367 (-) | 5851 | intergenic | Near Ysr105 |
| 757452 (-) | 3965 | intergenic | Near Ysr105 |
| 757520 (+) | 3791 | Intergenic | Near Ysr105 |
| 757978 (-) | 4206 | Intergenic | Near Ysr105 |
| 1732316 (-) | 7024 | YPTB1440 | Hypothetical protein |
| 2242048 (-) | 12513 | pseudogene | transposase (non-functional) |
| 2475581 (-) | 4932 | intergenic | Upstream of *hns* |
| 3138894 (+) | 1658 | intergenic | T6SS island |
| 4176313 (-) | 13980 | YPTB3506 | *sspA* stringent starvation protein |
| 4176973 (-) | 11465 | YPTB3506 | *sspA* stringent starvation protein |
| 4176992 (+) | 9682 | YPTB3506 | *sspA* stringent starvation protein |
| 4459942 (+) | 9948 | YPTB3758 | *igaA* prevents activation of RcsC |
| 4459955 (+) | 8919 | YPTB3758 | *igaA* prevents activation of RcsC |
| 4459959 (-) | 11687 | YPTB3758 | *igaA* prevents activation of RcsC |
| 4459976 (-) | 9543 | YPTB3758 | *igaA* prevents activation of RcsC |
| 4460097 (+) | 9138 | YPTB3758 | *igaA* prevents activation of RcsC |
| 4460363 (-) | 10221 | YPTB3758 | *igaA* prevents activation of RcsC |
| 4460771 (+) | 10589 | YPTB3758 | *igaA* prevents activation of RcsC |
| 4460832 (+) | 8849 | YPTB3758 | *igaA* prevents activation of RcsC |
| 4461288 (+) | 9740 | YPTB3758 | *igaA* prevents activation of RcsC |

**Supplementary Table 2.**
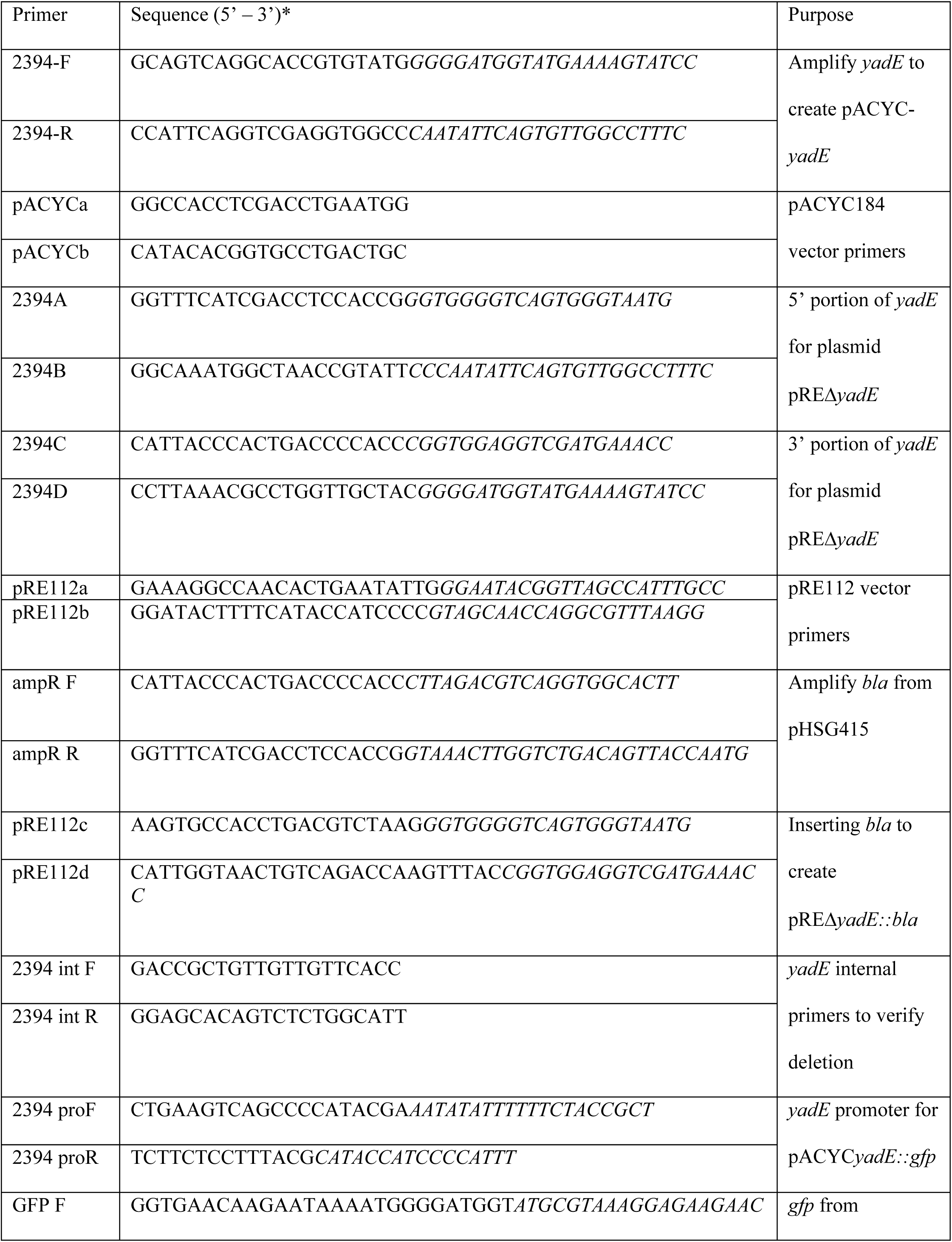
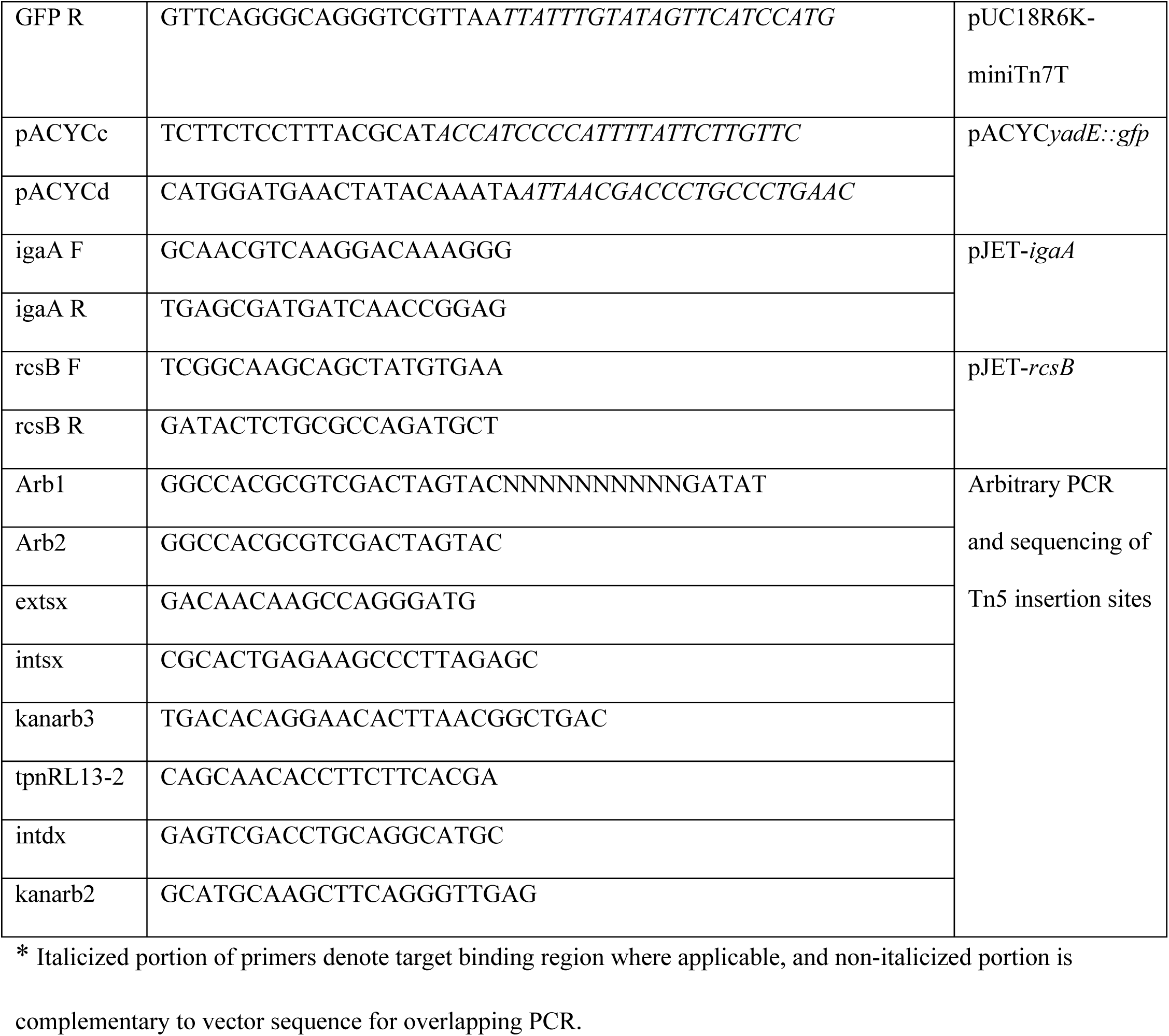
PCR Primers.

